# Towards increased shading potential: a combined phenotypic and genetic analysis of rice shoot architecture

**DOI:** 10.1101/2021.05.25.445664

**Authors:** Martina Huber, Magdalena M. Julkowska, Basten L. Snoek, Hans van Veen, Justine Toulotte, Virender Kumar, Kaisa Kajala, Rashmi Sasidharan, Ronald Pierik

**Author notes:** Author for correspondence: Ronald Pierik. **Author contributions** M.H., R.P. and R.S. designed the experiments, with additional input from K.K., J.T. and H.v.V.; M.H. performed all experiments, analysed the data, and wrote the article with contributions of all authors; M.M.J. carried out the haplotype analysis and assisted with statistical data analysis and data visualization; B.S. provided technical assistance for the genome-wide association studies and performed part of the analysis; H.v.V. provided assistance for statistical analysis; J.T. performed part of the experiment and measurements; V.K. contributed to research plan and experiment support at IRRI; R.P. serves as the author responsible for contact and ensures communication, supervised all experiments, revised the manuscript draft and together with R.S. conceived the research plan and project design.

## Abstract

Rice feeds more than half of the world’s human population. In modern rice farming, a major constraint for productivity is weed proliferation and the ecological impact of herbicide application. Increased weed competitiveness of commercial rice varieties requires enhanced shade casting to limit growth of shade-sensitive weeds and the need for herbicide. We aimed to identify traits that enhance rice shading capacity based on the canopy architecture and the underlying genetic components. We performed a phenotypic screen of a rice diversity panel comprised of 344 varieties, examining 13 canopy architecture traits linked with shading capacity in 4-week-old plants. The analysis revealed a vast range of phenotypic variation across the diversity panel. We used trait correlation and clustering to identify core traits that define shading capacity to be shoot area, number of leaves, culm and solidity (the compactness of the shoot). To simplify the complex canopy architecture, these traits were combined into a Shading Rank metric that is indicative of a plant’s ability to cast shade. Genome wide association study (GWAS) revealed genetic loci underlying canopy architecture traits, out of which five loci were substantially contributing to shading potential. Subsequent haplotype analysis further explored allelic variation and identified seven haplotypes associated with increased shading. Identification of traits contributing to shading capacity and underlying allelic variation presented in this study will serve future genomic assisted breeding programmes. The investigated diversity panel, including widely grown varieties, shows that there is big potential and genetic resources for improvement of elite breeding lines. Implementing increased shading in rice breeding will make its farming less dependent on herbicides and contribute towards more environmentally sustainable agriculture.

**One sentence summary:** Through screening a rice diversity panel for variation in shoot architecture, we identified traits corresponding to plant shading potential and their genetic constituents.

## Introduction

Rice feeds more than half of the world’s population as a staple food (Kennedy and Burlingame, 2003; Wing et al., 2018). In traditional rice farming, seedlings are transplanted into flooded paddy fields. This works as a natural way to prevent weed infestation, since it gives rice seedlings a size advantage in addition to flood-suppressed germination and growth of weeds. This practice is increasingly problematic, both because of the high manual labour input (Kumar and Ladha 2011; Chakraborty et al. 2017) and because global climate change is reducing the availability of fresh water not only for rice farmers but for the global agricultural sector (FAO, 2019; Oliver et al., 2019). Traditional rice farming system is transitioning towards direct-seeded rice, where rice seeds are directly sown into the fields. This practice drastically reduces the water requirement and labour input (Chauhan et al., 2017; Farooq et al., 2011; Kumar and Ladha, 2011). Besides all of its advantages, the major constraint for direct-seeded rice is abundant proliferation of weeds (Rao et al. 2017; Xu et al. 2019). In direct-seeded rice practice, rice seedlings are directly competing with weeds as they lose their seedling size advantage. Waterlogging cannot be applied to suppress emerging weeds, as most modern rice cultivars do not germinate under water (Chauhan, 2012; Ghosal et al., 2019; Kretzschmar et al., 2015). Currently, weeds are suppressed with herbicides, leading to evolution of herbicide-resistant weeds and ground water pollution. This creates a pressing need for deployment of sustainable weed management options (Chauhan, 2012a; Chauhan and Yadav, 2013; Mennan et al., 2012; Zhao et al., 2006a). One possible solution to this problem is to increase weed-competitiveness of the rice seedling (Rao et al., 2007; Sakamoto et al., 2006; Zhao et al., 2007).

Just like their wild ancestors, shade casting crop varieties compete with invading weeds by reducing the weed’s access to full sunlight, thereby impeding their growth. However, the traits contributing to shading potential were neglected or even selected against in breeding efforts, since tall plants and droopy leaves are generally considered as undesired, because it makes harvesting more difficult. Here we propose to develop weed-competitive rice varieties, by selecting for an ideotype with faster growth and high shade-casting potential on proximate weeds. Shoot architecture traits that help plants to gain advantage over their neighbours through light competition include: high number of leaves, increased tillering, large projected shoot area, increased planar angle of leaves and tillers (Andrew et al., 2015; Brainard et al., 2005; Mahajan and Chauhan, 2013; Seavers and Wright, 1999; Worthington and Reberg-Horton, 2013). Accelerated vertical growth might provide an additional advantage for outcompeting neighbours, yet plant height has been strongly selected against during green revolution of most cereals, including rice. Indeed, there exists great potential for weed suppression in cereal canopies, as has been shown for wheat, where a rapidly closing wheat crop canopy achieved through higher planting density, depleted weeds from access to light (Weiner et al., 2010).

Building on the idea to increase shading for improved weed competitiveness, here we examined the variation in rice shoot architecture, derived the traits that contribute to shading potential, and identified genetic loci associated with shading potential. The shading potential was defined here as high ground cover and early growth vigour. We determined key architectural characteristics of shading potential in the early growth phase. For this, (1) we phenotyped a rice diversity panel of 344 globally distributed varieties where we recorded 13 quantitative traits. Based on these, (2) we determined key architectural characteristics of shading potential in early growth phase. (3) We combined these core traits into one parameter to develop the Shading Rank, where all studied rice varieties were ranked for their shading potential. (4) Genome-wide association study (GWAS) revealed association with five genetic loci for traits contributing to shade potential. The results of this study form a primer to identification of alleles contributing to increased shading and early plant vigour.

## Results

### Shoot architecture variation between rice varieties

In order to establish the variation in shading potential, and resulting increased weed-competitiveness, within the rice diversity panel (Supplemental Table 1) we measured 13 traits on 4-week-old seedlings in the screenhouse (Figure 1, Table 1, Supplemental Table 2).

**Figure 1:**
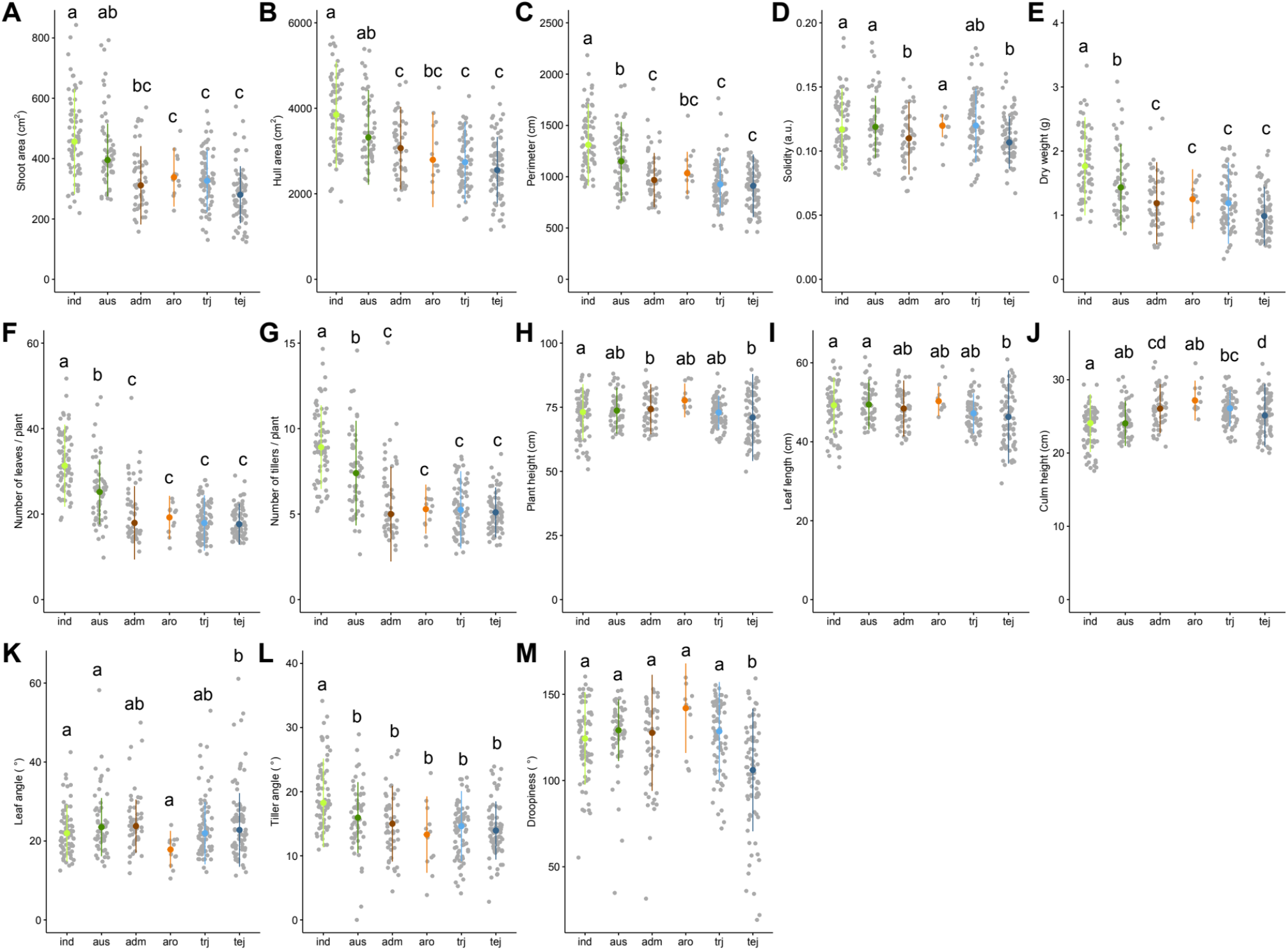
Shoot traits in rice differ between subpopulations. Distribution of investigated shoot traits in the screened diversity panel. The plots represent the trait value (y-axis) observed for varieties grouped according to different subpopulations on x-axis. A) Shoot area [cm2], B) Hull area [cm2], C) Perimeter [cm], D) Solidity, E) Dry weight [g], F) number of leaves, G) Number of tillers, H) Plant height [cm], I) Leaf length [cm], J) Culm height [cm], K) Leaf angle [°], L) Tiller angle [°] and M) Droopiness [°]. Each data point represents the mean out of 8 replicates for each of the 344 varieties. The colours represent different groups of subpopulations, ind – *indica*, aus, adm – *admixed*, aro -*aromatic*, trj – *tropical japonica* and tej – *temperate japonica*. The letters in the graphs represent the significantly different groups, as determined with Tukey’s HSD with p-value < 0.05. Mean values for all 13 traits and the sum of the normalized traits including results for Tukey’s pairwise post hoc test can be found in Supplemental Table 2.

**Table 1:**
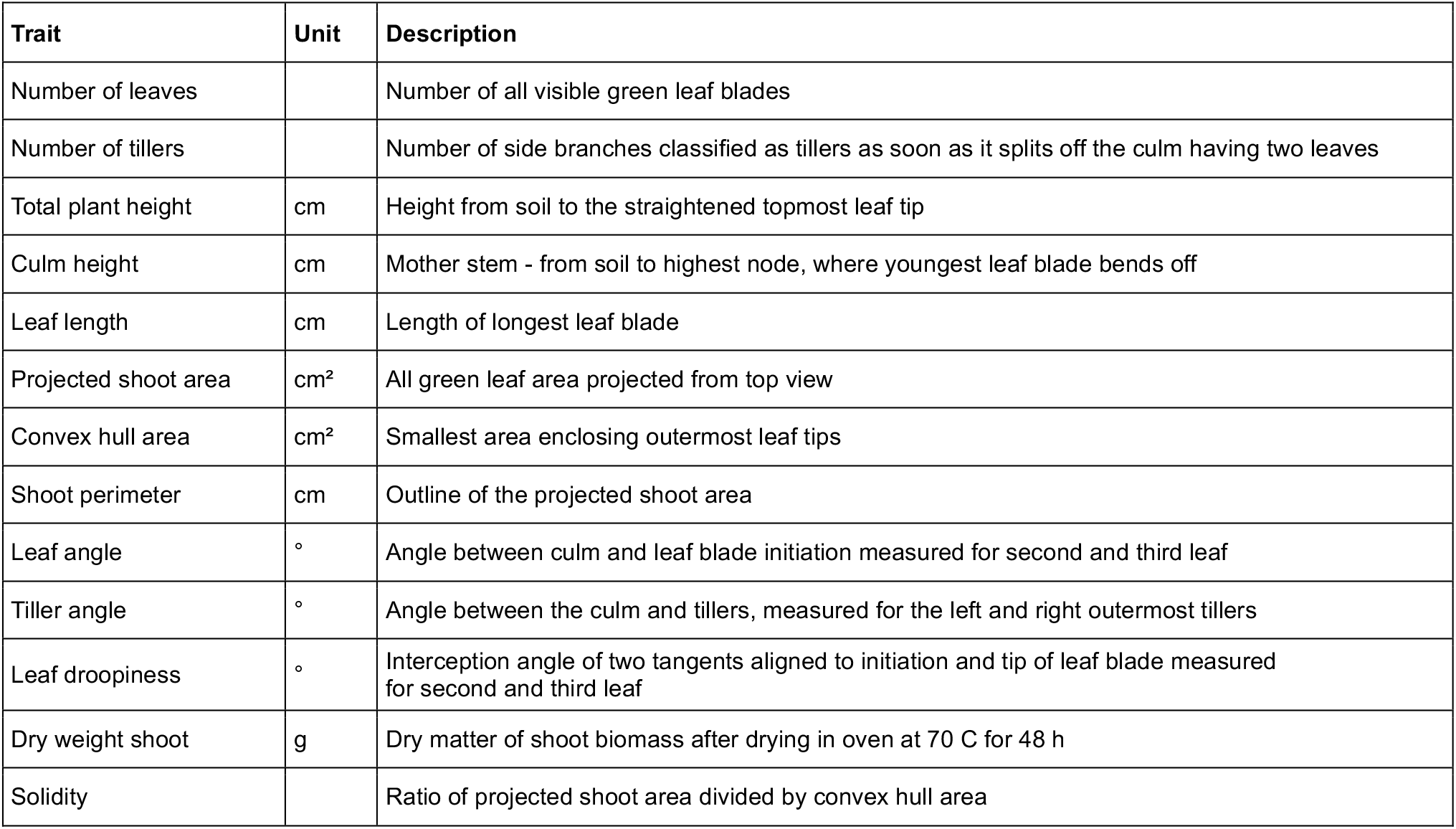
Description of 13 investigated shoot traits.

| Trait | Unit | Description |
| --- | --- | --- |
| Number of leaves |  | Number of all visible green leaf blades |
| Number of tillers |  | Number of side branches classified as tillers as soon as it splits off the culm having two leaves |
| Total plant height | cm | Height from soil to the straightened topmost leaf tip |
| Culm height | cm | Mother stem - from soil to highest node, where youngest leaf blade bends off |
| Leaf length | cm | Length of longest leaf blade |
| Projected shoot area | cm<sup>2</sup> | All green leaf area projected from top view |
| Convex hull area | cm<sup>2</sup> | Smallest area enclosing outermost leaf tips |
| Shoot perimeter | cm | Outline of the projected shoot area |
| Leaf angle | ° | Angle between culm and leaf blade initiation measured for second and third leaf |
| Tiller angle | ° | Angle between the culm and tillers, measured for the left and right outermost tillers |
| Leaf droopiness | ° | Interception angle of two tangents aligned to initiation and tip of leaf blade measured for second and third leaf |
| Dry weight shoot | g | Dry matter of shoot biomass after drying in oven at 70 C for 48 h |
| Solidity |  | Ratio of projected shoot area divided by convex hull area |

Substantial variation was observed for all measured traits among the varieties belonging to different subpopulations (Figure 1; Supplemental Table 2). The *indica* subpopulation showed highest dry weight, number of leaves, and number of tillers followed by *aus* subpopulation and *aromatic, tropical* and *temperate japonica* ranked lowest in these parameters (Supplemental Table 2). Shoot and hull area were also observed to be higher in *indica* and *aus* subpopulations, intermediate in *aromatic* subpopulation and lowest in *japonicas* and *admixture* subpopulations. *Indica* and *aus* on average develop the most compact shoots (highest solidity), contrasting with the low solidity of *japonicas* and *aromatic*. In plant height, *indica* lines and *temperate japonica* were shortest and *aromatic* subpopulation were tallest. When taking the entire diversity panel of 344 varieties, five traits (shoot area, hull area, solidity, plant height and dry weight) already showed a significant difference between the individual varieties at four weeks after sowing (Supplemental Table 2). When grouped together in subpopulations, all traits showed significant differences between subpopulations (Supplemental Table 2). Overall, it appears that relatively large variation between subpopulations was observed for traits related to area and branchiness related traits, whereas traits related to height showed only little variation between subpopulations. These differences are clearly determined by differences in genetic background since the growth conditions were constant. The high variation observed for traits related to shading potential suggests that the investigated rice diversity panel has sufficient variation to improve shading of the elite-breeding varieties.

### Correlation of shoot architectural traits

To explore the relationship between individually measured traits, and determine which traits are independent of each other, we performed a Pearson correlation analysis (Figure 2A, Supplemental Figure 1). Shoot area and hull area showed strong positive correlation with shoot dry weight. Leaf and tiller number were highly correlated with dry weight. Height-associated traits, such as plant height, culm height and leaf length, were positively correlated with each other. On the other hand, a negative correlation was found between culm height and number of leaves and tillers. Solidity, leaf angle, tiller angle and droopiness did not exhibit strong correlation with other measured traits.

**Figure 2:**
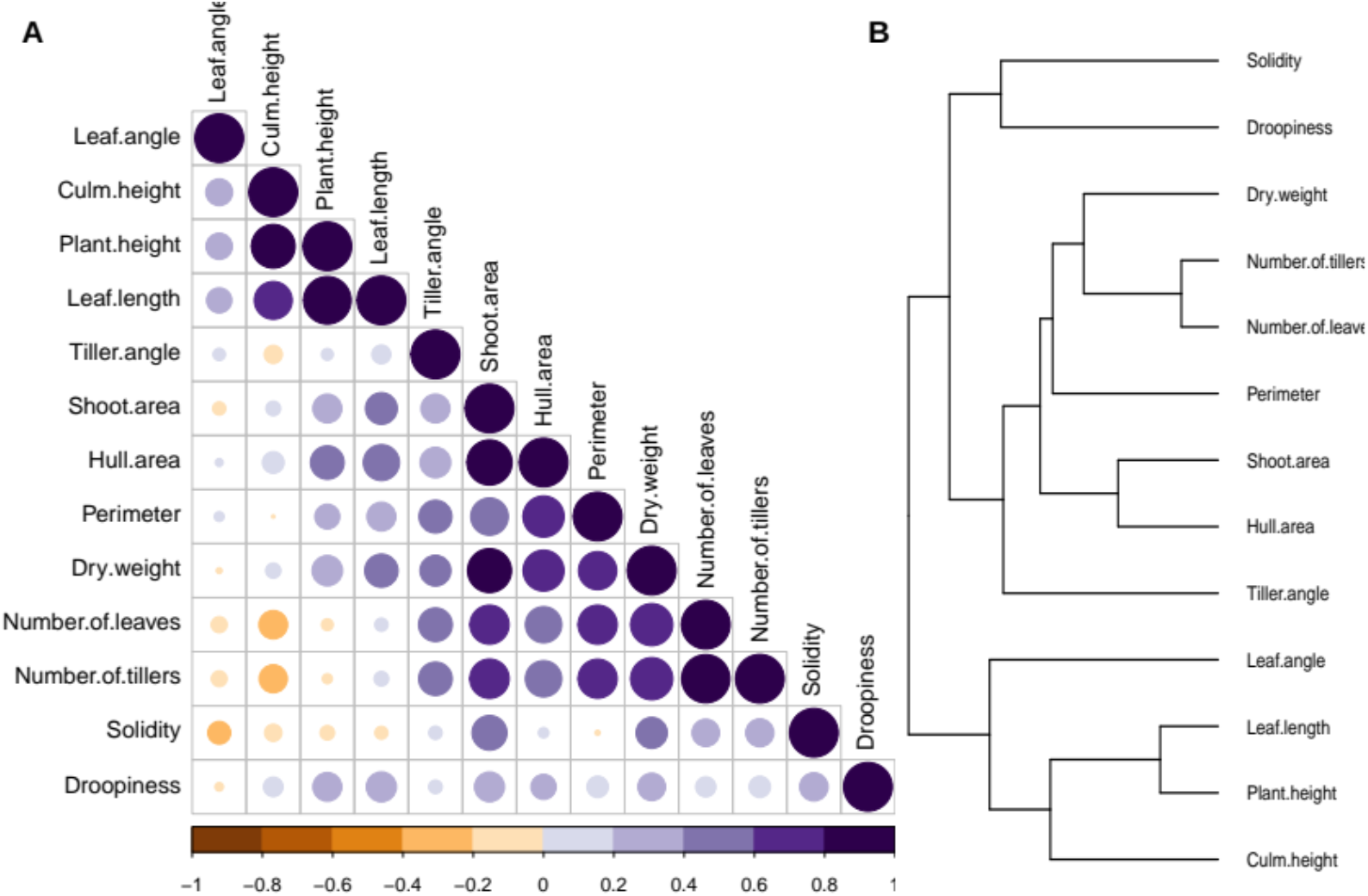
Correlation and clustering of 13 shoot traits defines core groups of traits. A) Pearson Correlation coefficients between traits. The colour and size of the circles reflect the strength of the correlation. B) Hierarchical Cluster Analysis. Traits are clustered using ward.D2 method. Rows represent 13 studied shoot traits. The values of individual samples are normalized per trait using z-Fisher transformation scaled prior to clustering. Based on a cut off at seven clusters and together with the correlation coefficients, we grouped together the traits into defined core groups.

To examine the types of canopy architecture exhibited within rice diversity panel, we performed hierarchical clustering (Figure 2B), resulting in seven trait clusters. The clustering shows how traits are grouped together according to the patterns observed across all rice varieties. Taking the correlation and clustering analyses together, we can determine core groups of traits: area-related (shoot area, hull area, perimeter), branchiness (number of leaves and tillers and dry weight), height-related (plant and culm height and leaf length), solidity, leaf angle, tiller angle and droopiness (Table 2).

**Table 2:** Core groups of shoot traits. For core groups with multiple traits, we have selected a representative trait as the core trait, shown in bold.

| Core groups | Measured shoot architectural traits |
| --- | --- |
| Area | **Projected shoot area**, convex hull area, perimeter |
| Branchiness | **Number of leaves**, number of tillers, dry weight |
| Height | **Culm height**, leaf length, plant height |
| Solidity | Solidity |
| Leaf angle | Leaf angle |
| Tiller angle | Tiller angle |
| Droopiness | Droopiness |

### Defining “shading potential”

The shading potential of a plant determines the effectiveness with which it can cover ground area. The shading potential increases with an increasing number of leaves and tillers (branchiness), the size of the leaf and tiller angles, and the shoot area. Additionally, the shading potential accounts for plant height, as it offers competitive advantage to shading shorter weed plants. Therefore, plants with an increased height, number of leaves and wider angles are considered more vigorous, and thus likely to outcompete weeds for sunlight by casting more shade. With the aim of finding the ideal plant with highest shading potential for effective weed competition, we need to determine varieties with high values for core traits. The distribution of the different varieties with respect to the core trait groups: area, branchiness, height and solidity are shown in Figure 3, together with top images of representative varieties.

**Figure 3:**
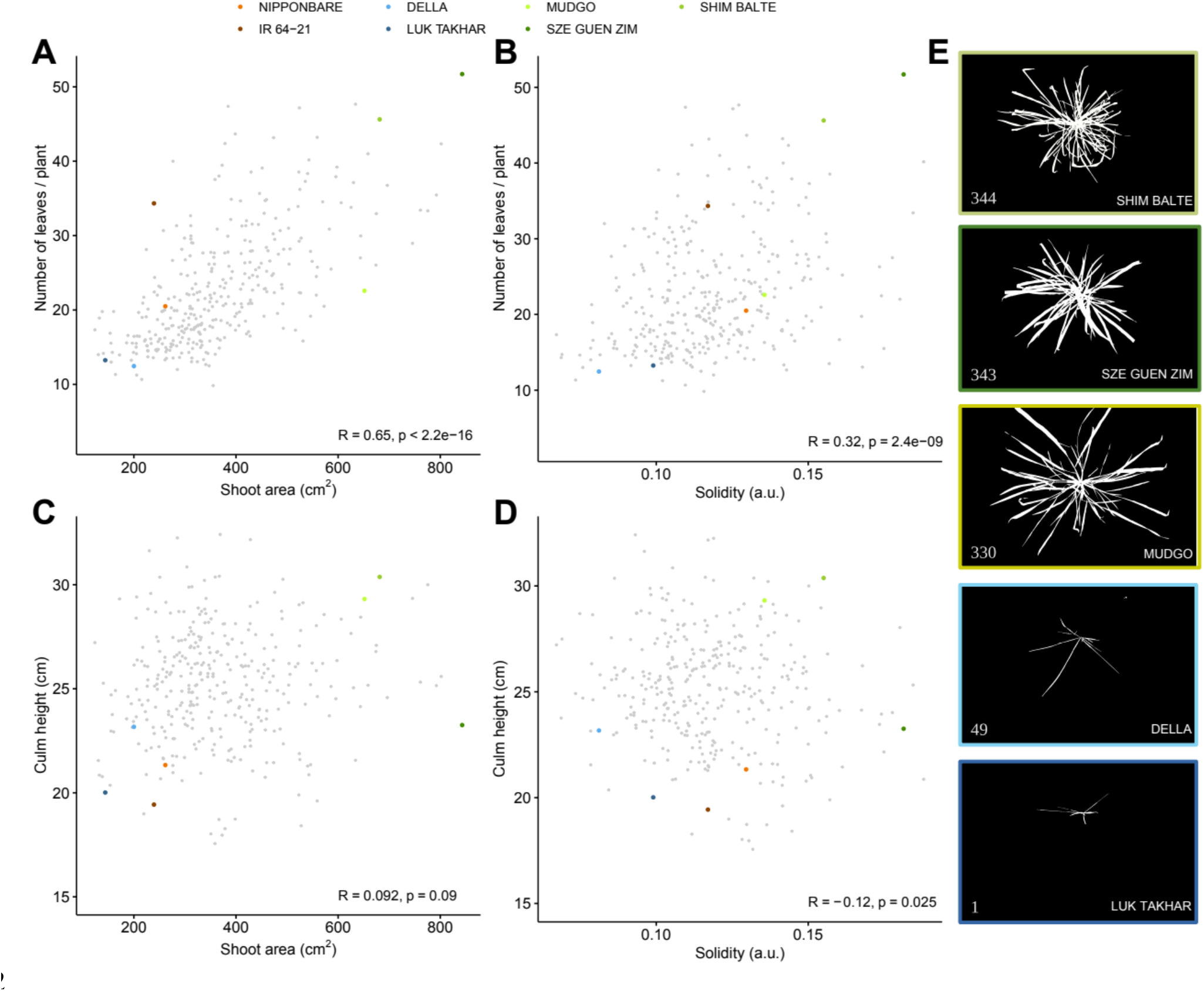
Visualization of shading potential in the investigated rice diversity panel based on cor traits for the Shading Rank. A) - D) Scatter plots showing the distribution of 344 rice varieties in pair-wise combination of four core traits, shoot area, number of leaves, solidity and culm height. Representative high (344, 343 and 330) and low (49 and 1) ranking varieties together with Nipponbare (73) and IR 64-21 (74) are highlighted in colours. B) Top view images of representative varieties, with colour coded frames. Numbers are respective Shading Ranks as found in Table 3.

To quantify shading potential, we ranked varieties for the sum of the core traits contributing to shading potential (projected shoot area, number of leaves, solidity, culm height, leaf angle, tiller angle and leaf droopiness, bold in Table 2). To account for the differences in measured units and unit ranges, for each trait, the values were rescaled to a range from 0 to 100, whilst keeping the relative differences of trait-values between different varieties unchanged and these relative differences of trait values are also reflected in the sum of the normalized trait values. Varieties then were ranked according to their sum of normalized trait values, from 344 (highest) to 1 (lowest), resulting in the Shading Rank (for detailed information see Methods section - Data processing and statistical analysis).

The Shading Rank gives a quantitative measure of the shading potential for a certain variety and indicates where a specific variety ranks with respect to the entire diversity panel (Supplemental Table 3). Although this ranking allows insight into the distribution of shading potential and the identification of expected strong and weak shaders, a limitation of this ranking is that it applies only within the diversity panel tested. Shoot size is one of the major factors contributing to overall shading potential. Since the diversity panel was evaluated 28 days after sowing, the large shoot size of high-ranking varieties also indicates faster growth and seedling vigour. The varieties with the highest Shading Rank were Shim Balte, Sze Guen Zim, Paraiba Chines Nova, P 737, Shirkati and Sabharaj, while varieties with lowest Shading Rank were Luk Takhar, Guineandao, Bul Zo and Shirogane. From the 25 highest ranking varieties, 14 belong to the *indica subpopulation* and eight to *aus*. Low scoring varieties in terms of shading potential include widely-grown varieties such as IR 64 and Nipponbare, ranking 74^th^ and 73^rd^ respectively (Table 3, Figure 3). This suggests that some of the current elite rice varieties could have a rather poor shading potential, and through breeding with varieties from *indica* and *aus* subpopulations, the shading potential and weed-competitiveness can possibly be increased.

**Table 3:** Shading Rank for ten highest and ten lowest ranking varieties, and for varieties of special interest (Mudgo, IR 64-21, Nipponbare and Della) with normalized core trait values (between 0 as lowest and 100 highest) compared to the min and max values within the screened panel and the sum of the core traits. Varieties in bold are visualized in Figure 3. The Shading Rank ranges from 344 as the highest and 1 as the lowest. The list of Shading Ranks for the entire panel can be found in Supplemental Table 3.

| Variety | Subpopulation | Shoot area.norm | Number of leaves.norm | Solidity.norm | Culm height.norm | Leaf angle.norm | Tiller angle.norm | Droopiness.norm | SUM_norm_traits | Shading Rank |
| --- | --- | --- | --- | --- | --- | --- | --- | --- | --- | --- |
| **SHIM BALTE** | **aus** | **78** | **85** | **73** | **86** | **94** | **65** | **79** | **561** | **344** |
| **SZE GUEN ZIM** | **ind** | **100** | **100** | **95** | **38** | **15** | **55** | **67** | **470** | **343** |
| PARAIBA CHINES NOVA | ind | 77 | 55 | 51 | 64 | 25 | 100 | 90 | 462 | 342 |
| P 737 | aus | 91 | 56 | 69 | 84 | 42 | 49 | 68 | 458 | 341 |
| SHIRKATI | aus | 93 | 61 | 68 | 51 | 8 | 85 | 80 | 446 | 340 |
| SABHARAJ | ind | 94 | 78 | 63 | 54 | 23 | 57 | 73 | 443 | 339 |
| PAUNG MALAUNG | aus | 89 | 56 | 97 | 52 | 16 | 45 | 85 | 440 | 338 |
| NIRA | ind | 80 | 64 | 56 | 47 | 32 | 70 | 82 | 431 | 337 |
| SATHI | aus | 67 | 59 | 66 | 73 | 22 | 52 | 81 | 420 | 336 |
| MTU9 | ind | 86 | 46 | 57 | 79 | 19 | 48 | 82 | 417 | 335 |
| **MUDGO** | **ind** | **73** | **30** | **57** | **79** | **20** | **53** | **95** | **407** | **330** |
| **IR 64-21** | **ind** | **16** | **59** | **41** | **13** | **16** | **32** | **78** | **254** | **74** |
| **NIPPONBARE** | **tej** | **19** | **25** | **52** | **25** | **13** | **42** | **77** | **253** | **73** |
| **DELLA** | **trj** | **11** | **6** | **12** | **38** | **66** | **46** | **56** | **234** | **49** |
| COCODRIE | trj | 10 | 11 | 22 | 39 | 23 | 26 | 38 | 168 | 10 |
| L 202 | trj | 1 | 10 | 9 | 27 | 14 | 44 | 61 | 166 | 9 |
| TRIOMPHE DU MAROC | tej | 2 | 10 | 51 | 52 | 22 | 25 | 2 | 165 | 8 |
| S 4542 A 3-49B-2-12 | trj | 4 | 8 | 7 | 48 | 5 | 43 | 43 | 159 | 7 |
| TAINAN IKU 487 | tej | 5 | 24 | 38 | 36 | 12 | 19 | 19 | 154 | 6 |
| PI 298967-1 | adm | 5 | 11 | 1 | 42 | 17 | 34 | 34 | 143 | 5 |
| SHIROGANE | tej | 4 | 17 | 14 | 19 | 12 | 34 | 43 | 142 | 4 |
| BUL ZO | tej | 10 | 8 | 20 | 45 | 22 | 21 | 11 | 137 | 3 |
| GUINEANDAO | adm | 10 | 14 | 9 | 38 | 8 | 40 | 9 | 127 | 2 |
| **LUK TAKHAR** | **tej** | **3** | **8** | **26** | **17** | **5** | **44** | **0** | **103** | **1** |

Interestingly, none of the top-ranking varieties showed the highest values for all core shading traits (Figure 3). For example, Sze Guen Zim ranks highest for shoot area and number of leaves, but is one of the lower-ranking varieties for culm height. The accession with the highest Shade Ranking (344), Shim Balte has a very high number of leaves and solidity, but has a close to average culm height. Mudgo reaches a rank of 340, despite its relatively low number of leaves and solidity. Della, a variety with a low rank of 49, ranks low for all traits except for culm height. Luk Takhar is at the bottom end of the ranking and shows low values for all core traits. The core traits that determine shading potential: shoot area, number of leaves, solidity and culm height are only weakly correlated (Figure 2, Figure 3), illustrating the diverse strategies to reach high shading potential. It is therefore important to include all of the four core traits, in addition to the angle related traits, for a comprehensive evaluation of shading potential.

### Predicted competitive varieties are casting more shade

To validate our Shading Rank and assess functional shading capacity, we grew varieties with varying Shading Rank and evaluated them for canopy shading. We selected two of the predicted competitive (Shim Balte with a Shading Rank of 344 and Mudgo ranking 330) and two predicted non-competitive rice varieties (Luk Takhar ranking 1 and Della ranking 49) (Figure 3, Table 3). By measuring the light quantity under the canopies of selected varieties (Supplemental Table 4), we indeed observed strong shading by varieties with a high Shading Rank (Shim Balte and Mudgo) and low shading by varieties with a low Shading Rank (Luk Takhar and Della). This result validates our Shading Rank, at least for the varieties tested and the selection of shoot architecture traits to effectively predict shade casting.

### SNPs associated with seedling establishment and shoot architectural traits

The high phenotypic variability found in the studied diversity panel (Supplemental Table 5), together with the high genetic variation (Wang et al., 2018b) provides a strong basis for a GWAS. We observed high narrow-sense heritability for all measured traits (Supplemental Table 6). We investigated the genomic trait associations on two different SNP sets, both with two different software packages (lme4QTL (Ziyatdinov et al., 2018) and Genomic Association and Prediction Integrated Tool (GAPIT) (Tang et al., 2016; Wang et al., 2018c), see methods for detailed description). The total list of p-values for SNPs association across all measured traits can be found at https://doi.org/10.5281/zenodo.4730232 (Supplemental Data 3).

Three genomic regions were associated with plant height located on chromosome 3, 5 and 6 (Figure 5). The peak on chromosome 3 was also detected for other height related traits: culm height and leaf length (Supplemental Data 4). Overall, the associations with culm height showed lower LOD scores (Supplemental Data 4). The results for tiller angle and droopiness reveal strong associations with SNPs on chromosome 1 and chromosomes 1 and 10, respectively (Supplemental Data 4). Despite solidity being a very complex and likely a poly-genic trait, the analysis revealed a strong association with 14 SNPs in the locus on chromosome 3 (Figure 5). The associations between leaf or tiller number, found for SNPs on chromosomes 11 and 12, were shared between these two traits (Supplemental Data 4). These two loci were also found for dry weight. This suggests that the genetic components underlying formation of new leaves and tillers might have a common genetic constituent, consistent with high correlation in their phenotypes (Figure 2). The analysis for dry weight revealed significant associations on chromosomes 3, 7 and 12, overlapping with the associations found for shoot area (Figure 5). When taking together shading potential as the sum of all core traits, a GWAS on this composite trait yielded a rather random pattern of SNP associations (Supplement Figure 4). This further highlights our earlier findings (Figure 4), that shading can be achieved through various strategies and shading potential, as such, is genetically a highly complex trait. Therefore, genetic mapping of shoot architecture components that contribute to shading capacity is much more effective approach in identifying genetic components that contribute to shading and potential weed competitiveness.

**Figure 4:**
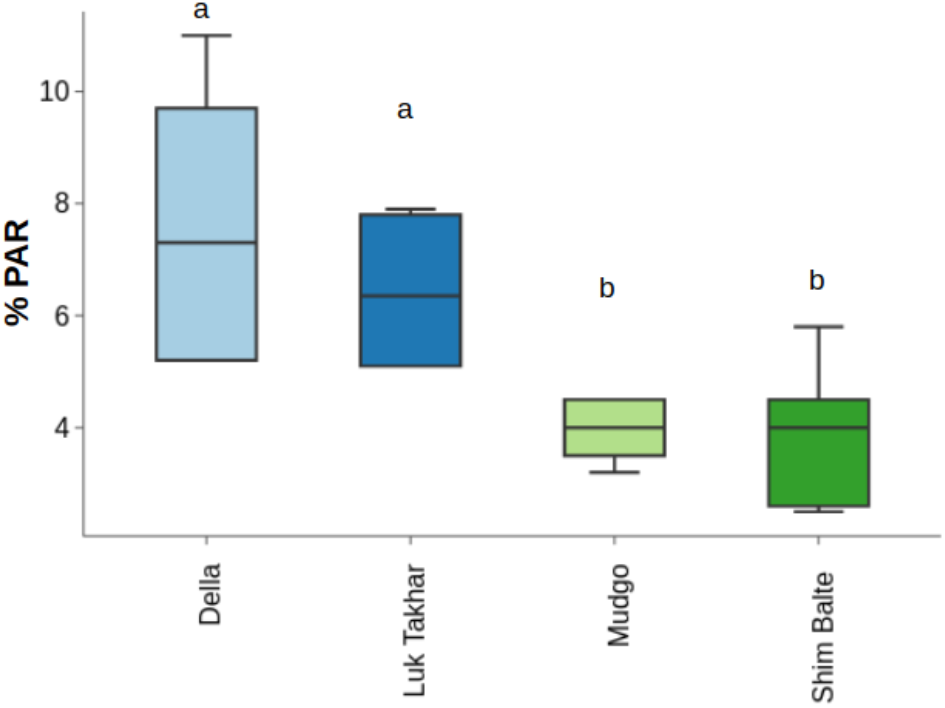
Shading Rank predicts the canopy shading capacity of high and low ranking rice varieties. Significant difference in shading capacity between canopies of different rice varieties at five weeks after sowing. The plot shows the reduction in light intensity (% PAR) measured at the ground level under the rice canopy compared to above the canopy, for different rice varieties on x-axis, where Della and Luk Takhar were classified as non-competitive (blue) with Shading ranks of 49 and 1, respectively and Mudgo and Shim Balte as competitive (green) with Shading Ranks of 330 and 344, respectively. Letters indicate significance (ANOVA with Tukey’s pairwise comparison post hoc test p < 0.05). Measured PAR values (photosynthetic active radiation of 400-700 nm waveband) can be found in Supplemental Table 4.

**Figure 5:**
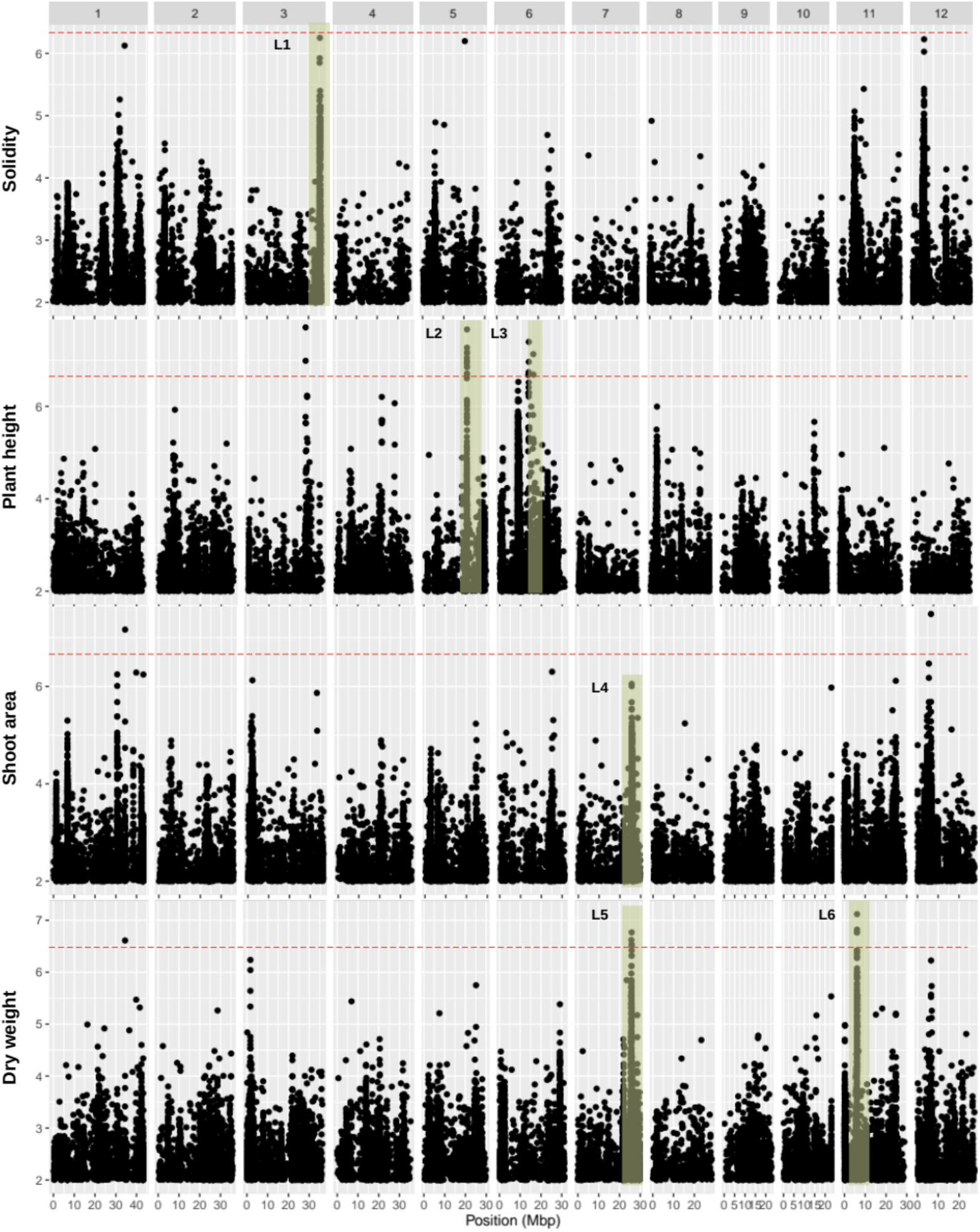
GWAS identifies putative the genetic regions underlying shoot architectural traits and seedling vigour in 4-week-old rice seedlings, reflecting the early vegetative growth stage. We used single-trait genome-wide association studies (GWAS) with a mixed linear model (MLM) for plant height, solidity, shoot area and dry weight. The Manhattan plots depict the single nucleotide polymorphisms (SNPs) with minor allele frequencies (MAF) > 0.05. Negative logarithmic p-values on the y-axis, for 1.7 M SNPs across the 12 rice chromosomes along the x‐axis. Dashed red lines indicate significance threshold set at –log10(p-value) > 7.5. Genomic regions highlighted in green are loci of interest (numbered L1 – L6).

### Identification of alleles associated with increased sharing potential

The genomic regions that consisted of multiple SNPs above the Bonferroni threshold within the calculated local average LD (Table 5) were investigated in more detail. Since the traits related to the canopy shading potential are the primary focus of this work, we prioritized the loci associated with culm height, shoot area, solidity and dry weight. Locus 4 of shoot area overlaps with locus 5 detected for dry weight (Figure 5, Table 5) and was therefore taken together in the follow up analysis. In total we determined five loci to be followed up with a haplotype analysis to identify specific alleles which could contribute to shading potential. By grouping varieties according to SNPs within one coding region, and examining the differences between identified haplotypes, we identified allelic variation associated with high shading potential (Figure 6, Table 6).

**Table 4:**
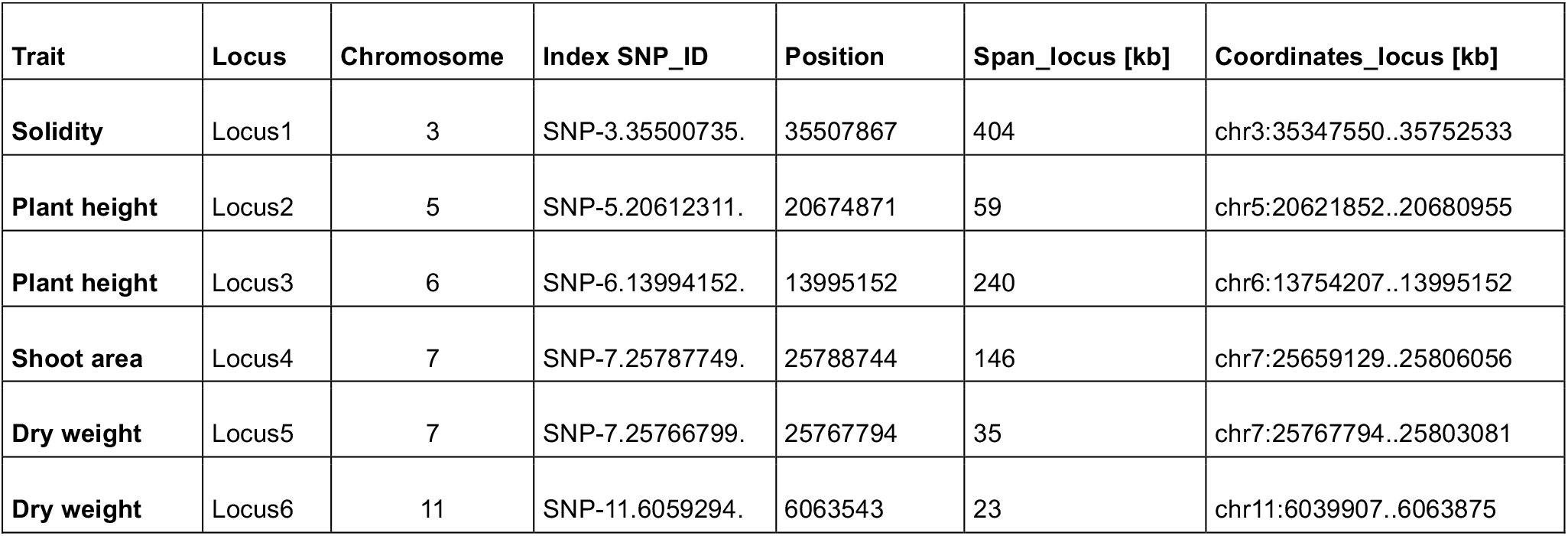
Loci of interest for traits of core groups for shading potential. (solidity, plant height, shoot area, and dry weight) with significant SNPs (LOD > 5 as index SNPs) and clumped SNPs (LOD > 4) in local LD up- and downstream. Full list of SNP positions in loci of interest can be found in Supplemental Table 7.

**Table 5:** Summary of determined loci of interest with the Locus ID and gene annotation. Loci represented in Figure 6 are highlighted in bold. Full list of SNP positions in loci of interest with gene annotation and gene ontology categories can be found in Supplemental Table 7.

| Trait | Locus | Chromosome | Locus_ID | Gene annotation |
| --- | --- | --- | --- | --- |
| Solidity | Locus1 | 3 | Os03g0841800 | GSK3/SHAGGY-like kinase |
|  |  |  | Os03g0841850 | Hypothetical protein. |
|  |  |  | Os03g0843700 | FAR1 domain containing protein. |
|  |  |  | **Os03g0845000** | **Similar to Pirin-like protein.** |
|  |  |  | **Os03g0845700** | **Similar to RPB17 (Fragment).** |
|  |  |  | Os03g0845800 | Conserved hypothetical protein. |
|  |  |  | Os03g0848700 | Coiled-coil, nucleotide-binding, and leucine-rich repeat protein |
| Plant height | Locus2 | 5 | Os05g0420500 | Conserved hypothetical protein. |
|  |  |  | **Os05g0420600** | **Cytochrome c.** |
|  |  |  | **Os05g0420900** | **Conserved hypothetical protein.** |
| Plant height | Locus3 | 6 | Os06g0269300 | TolB-like domain containing protein. |
|  |  |  | Os06g0346300 | acyl-CoA oxidase/ oxidoreductase |
| Shoot area | Locus4 | 7 | **Os07g0623200** | **Heavy metal transporter protein; ATPase, P-type.** |
|  |  |  | Os07g0623501 | Hypothetical gene. |
|  |  |  | Os07g0623600 | Similar to mRNA, clone: RTFL01-43-H20. |
| Dry weight | Locus5 | 7 | Os07g0623200 | Heavy metal transporter protein; ATPase, P-type. |
|  |  |  | Os07g0623501 | Hypothetical gene. |
|  |  |  | Os07g0623600 | Similar to mRNA, clone: RTFL01-43-H20. |
| Dry weight | Locus6 | 11 | **Os11g0216000** | **Pyruvate kinase family protein.** |

**Figure 6:**
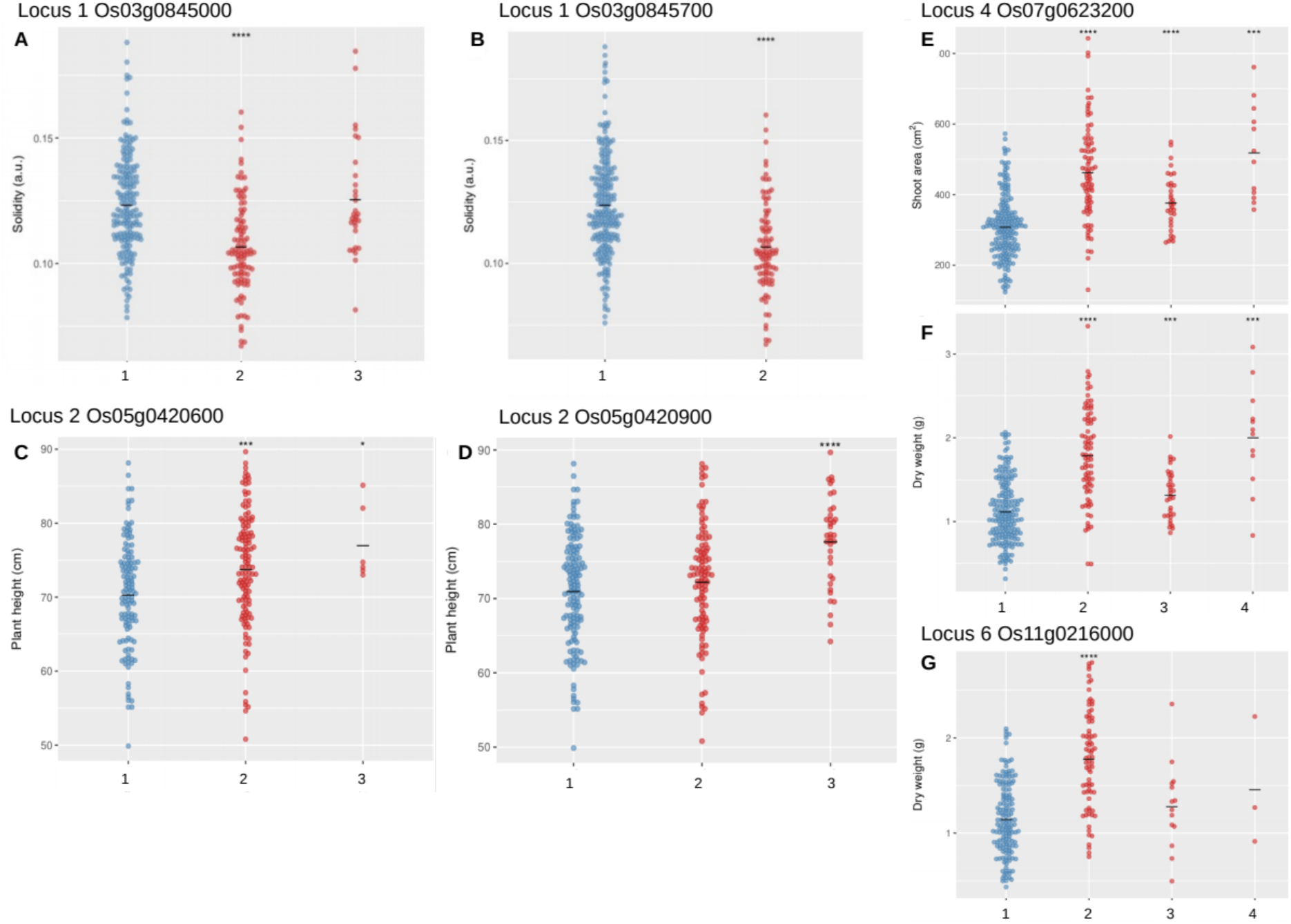
Haplotypes for genes of interest associated with increased trait values. Locus 1 was detected for solidity with haplotypes in the coding sequence of the genes A) Os03g0845000 consisting of two SNPs and B) Os03g0845700 consisting of one SNPs. Locus 2 was detected for plant height with haplotypes in the coding sequence of the genes C) Os05g0420600 consisting of four SNPs and B) Os05g0420900 consisting of six SNPs. Locus 4 was detected for shoot area and dry weight with haplotypes in the coding sequence of the gene Os07g0623200 consisting of four SNPs shown for E) shoot area and F) dry weight. Locus 6 was detected for dry weight encoding only one gene G) Os11g0216000 with haplotypes consisting of nine SNPs. Dot plots for t-test, comparing each haplotype with the most abundant (blue) haplotype, on core traits for shading potential. Y-axis trait value, x-axis groups of haplotypes. Additional information about the detected genes can be found in Table 5 and dot plots for haplotypes for all 13 traits found in loci of interest are shown in Supplemental Figure 5.

The haplotypes of two coding regions in locus 1 (Figure 6 A-B), associated with solidity, were observed to have significantly lower solidity than the most abundant haplotype identified for the respective coding regions. These are annotated as Os03g0845000 (Pirin-like protein) and Os03g0845700 (similar to RPB17 fragment). Haplotypes of two coding regions in locus 2 (Figure 6 C-D), associated with plant height, Os05g0420600 (Cytochrome C) and Os05g0420900 (conserved hypothetical protein), contained taller plants than the most abundant haplotype. In locus 4, associated with shoot area and dry weight, we found that only one gene (Os07g0623200, ATPase and heavy metal transporter protein) showed clear separation across the haplotypes, where all the non-reference haplotypes showed higher shading potential, indicated by higher shoot area and dry weight (Figure 6 E-F). For locus 6, associated with dry weight, we found only one gene Os11g0216000 encoding Pyruvate kinase family protein, we found that the second most abundant haplotype was associated with increased shading due to higher dry weight of varieties that were sharing this specific combination of SNPs.

## Discussion

We studied phenotypic and genetic variation in rice shoot architecture to identify traits and their underlying genetic loci that contribute to canopy shading. We investigated variability across a natural rice diversity panel in shoot architecture at the early vegetative stage. The traits investigated here encompass both early vigour and shade casting through shoot architecture, which are hypothesized to contribute to weed suppression in rice fields. Traits related to shoot architecture, such as leaf angle or droopiness, are of special interest as they do not require substantial resource investment while creating more optimal 3D distribution of the shoot biomass for an increased shading potential. Other traits, such as leaf area, number of leaves or shoot biomass, likely require considerable resource investments and are typically associated with growth vigour i.e. rapid seedling establishment.

In our screen for variation in shoot architecture traits we found significant differences between subpopulations, where varieties with an *indica* background have highest shading potential and *temperate japonica* the least. We found *admixed, tropical japonica* and *aus* subpopulations to typically range between *temperate japonica* and *indica*. This pattern could be found in the majority of the measured traits and is in line with the phylogenetic relatedness of the different subpopulations (Eizenga et al., 2014; Liakat Ali et al., 2011; McCouch et al., 2016; Zhao et al., 2011). This indicates that phylogenetic relatedness is an important component that determines phenotypic variation in shoot architecture and shading potential.

### Identification of core shading traits through correlation analysis

In order to summarize the information contained in all the investigated traits into one parameter indicative for the shading potential, we performed an extensive correlation analysis. By assessing the correlation between individual traits, we identified how all measured traits are related to one another and identified core traits that capture the observed variance (Figure 2). We identified groups of traits related to branchiness (number of leaves and tillers) and height (plant height, culm height and leaf length). The correlations between traits encapsulated within a trait group simply underlines the natural growth pattern; the more tillers a plant has, the more leaves it will have since each tiller will develop a certain number of leaves. Strong correlation was previously observed between tiller formation and relative growth rate (Dingkuhn et al., 2001). Likewise, in our study number of leaves and leaf area were positively correlated with shoot dry weight (Figure 2, Supplemental Figure 1). This well-established relationship (Caton et al., 2003; Dingkuhn et al., 2001; Poorter et al., 2012) can be explained due to a larger shoot area providing higher capacity for photosynthesis and thereby leading to higher overall growth rate (Caton et al., 2003). Not all traits showed expected correlations. It could for example be assumed that an increased inclination angle of the leaf blade would make a leaf droopier. In fact, leaf angle appeared to be unrelated to leaf droopiness, whereas leaf length appeared to be positively correlated with droopiness. While solidity is the ratio of shoot area and hull area, it is only weakly correlated with shoot area (Figure 2, Supplemental Figure 1). This suggests that shoot solidity is independent of how large its total shoot area, leaf number or dry weight are. Since solidity indicates the uniformity of the plant’s ability to shade its circumference, it is a valuable trait for shading capacity analysis. Inverse correlations were found between branchiness (number of leaves and tillers) and height traits. This trade-off between height and branching is well-documented as apical dominance where height growth of the main shoot is promoted at the expense of branching (Roig-Villanova and Martínez-García, 2016; Teichmann and Muhr, 2015). Summarizing, the trends observed within this study are in line with earlier observations, whereas we identify new, informative trait groups that contribute independently to the shading potential of rice plants.

### Shading rank as a measure for shading potential

Shading potential can be defined in two-dimensional measures, such as ground cover or projected shoot area, or including a third dimension, where plant height is considered as space resource utilization (Zhang et al., 2019). We hypothesized that not only projected shoot area, but also solidity and height of the shoot are crucial for shading potential. For example, a large projected shoot area with low solidity would still leave many open spaces within a single plant’s sphere for light penetration where weeds can proliferate. Or the reverse, a very solid projected shoot area of one plant that does not extend very far, is likely to leave open spaces between crop plants where weeds could grow. It is, therefore, clear that an optimal combination of shoot architecture traits is needed for maximal shading and weed suppression (Figure 3, Table 3). Architecture traits that are associated with weed-competitiveness include leaf area, ground cover, specific leaf area, leaf area index, leaf angle, droopiness, tillering capacity and plant height (Caton et al., 2003; Dingkuhn et al., 2001; Haefele et al., 2004; Mahajan and Chauhan, 2013; Mennan et al., 2012; Namuco et al., 2009; Rao et al., 2007; Worthington and Reberg-Horton, 2013; Zhao et al., 2006b, 2007). In addition, plant biomass and early vigour are advantageous for competition against weeds (Haefele et al., 2004; Mahajan and Chauhan, 2013; Namuco et al., 2009; Worthington and Reberg-Horton, 2013; Zhao et al., 2006a), but these are not specific architecture traits.

To predict which components best describe a plant’s shading potential, we categorized the different traits into core groups of similarly behaving traits. We developed the Shading Rank, as a parameter that combines branchiness, solidity and height and leaf and tiller angles and droopiness. The varieties with highest shading potential belong to the *indica* and *aus* subpopulation, which have also been found in earlier studies to have higher yield and less weed biomass in weedy fields compared to *japonicas* (Zhao et al., 2006b). We propose that varieties that have a high Shading Rank, are likely the most weed-competitive varieties, whereas those that rank low are likely to be weak competitors. Indeed, our experiment proved that canopies of high-ranking varieties allow significantly less light penetration than low ranking ones (Figure 4). Interestingly, none of the investigated varieties resembled the full ideotype of a strongly shading plant according to the traits we examined (Figure 3), indicating there is substantial room for improvement. Early seedling vigour is particularly important for weed-competition during the critical period of weed control and some of high ranking varieties, such as Shim Balte, Paung Malaung and Sabharaj are also known by breeders for their early vigour. Increased shading ability is intrinsic to early vigour since it follows to some extent from large size. However, the Shading Rank proposed here is more comprehensive to additional traits such as solidity and plant architecture that may involve less resource investment than vigour traits. With this improved way of ranking a plant’s shading capacity, our study exemplifies a new method of selection for high-shading varieties and genetic loci associated with high-shade canopy architecture.

### Elucidating the genetic components of shading potential

#### Architecture

The SNP dataset from the rice diversity panel (Eizenga et al., 2014) was combined with the observed phenotypic variation to identify putative genetic loci underlying high shading potential. This variation (Figure 1, Supplemental Table 5) together with a high trait heritability (Supplemental Table 6) provides a strong basis for GWAS. Plant height and leaf length were associated with loci on chromosomes 5 and 6. The locus on chromosome 5 harbours two genes encoding Cytochrome C and a conserved hypothetical protein. The haplotype analysis revealed one allele for both genes that was associated with a highly significant increase in plant height. (Figure 4). The locus on chromosome 6 encodes the *Heading Date* (*Hd1*) locus that was also previously associated with plant height in vegetative rice plants (Zhang et al., 2012; Yang et al., 2014). Subedi et al. (2019) performed a GWAS on plant height at plant maturity and found peaks on chromosome 1 and 11, which could indicate that at different developmental stages plant height is determined by different genomic regions. However, Subedi et al (2019) used a specifically constructed genetic population stemming from six parents and this could explain why very different loci were identified. Interestingly, haplotypes associated with high culm height exhibit low plant height and vice versa (Supplemental Data 7). Haplotypes associated with high plant height are typically showing longer leaf length (Supplemental Data 7). While all the height related traits were highly correlated at phenotypic level (Figure 2), the lack of common loci for all the traits (Supplemental Data 4), and opposite trends within the haplotype groups (Supplemental Data 7) suggest that the three components of plant height are regulated independently at the genetic level.

We also report unique loci specific for solidity and for height related traits. We revealed one strong locus, with several significant SNP associations, on chromosome 3 for solidity (Figure 5). We propose that solidity, as mentioned previously, is an important shoot trait to take into consideration for weed-competitiveness, since high crop plant solidity likely indicates low potential for weeds to proliferate within the sphere of influence of crop individuals. It is surprising to find a single locus, uniquely associated with this complex trait. However, when we grouped varieties into haplotype groups for two coding regions (Os03g0845000 and Os03g0845700, Figure 6 A – B), encoding a Pirin-like protein and an RPB17 (Fragment) within this locus, the phenotype of the haplotype groups appeared to differ not just in solidity, but also shoot area, dry weight and leaf number (Figure 6 A-B, Supplemental Data 7).

In this analysis, we identified new genetic components of shading potential based on shoot architecture, and the alleles that might contribute to increased shade casting ability.

#### Vigour

Vigour-related traits (i.e., dry weight, shoot area, number of leaves) are all strongly correlated and share associated loci on chromosome 7, 11 and 12 (Figure 5, Supplemental Data 4). The locus on chromosome 11 was also reported by (Yang et al., 2014) for dry weight and fresh weight at the late tillering stage, which is comparable to the developmental stage studied here. A closer look at the locus found for dry weight on chromosome 11 revealed only one gene is located within the linkage disequilibrium of associated SNPs. Interestingly, the haplotype analysis for SNPs within Os11g021600, encoding a Pyruvate kinase family protein, revealed significant difference in dry weight between the two haplotype groups (Figure 6 G). The significant differences were also observed for shoot area and number of leaves and tillers. As only one gene was located within this locus and one specific haplotype was related with high biomass, this locus is a promising candidate for follow-up studies and promising to be included in breeding programmes. The locus on chromosome 7 associated with shoot area and dry weight (Figure 6 E and F), harbours two genes, where we found that the haplotypes were associated with an increased shoot area and dry weight but also increased number of leaves and tillers. QTLs for height at 7 and 14 days after sowing and fresh weight, in a study that involved exclusively *temperate japonica* genotypes (Cordero-Lara et al., 2016) were entirely non-overlapping with the loci identified here for these traits. This is most likely because of the different genetic make-up of the populations used, which inevitably leads to variation. Even though the GWAS results for number of leaves and dry weight revealed different genetic associations for each of these traits, the identified haplotypes affected both these traits in a similar way. The haplotypes associated with high projected shoot area also showed increased branchiness and dry weight (Supplemental Data 4). This might suggest that by selecting for a genetic locus associated with branchiness, the other traits contributing to shading potential might also be affected. This relationship is to be further studied in future reverse-genetic studies that could explore the role of identified candidate loci in increased shading potential as well as weed-competitiveness.

It should be kept in mind that rice is known to be a highly plastic species and we have performed our experiments under stable conditions in a controlled environment. In order to further translate our findings, and implement them in breeding programmes, it will be relevant to factor in architectural plasticity under field conditions. One obvious factor affecting architecture would be planting density and the associated changes in light composition and availability. Another so far neglected aspect of weed-competitiveness would be the root systems, for which the rapidly evolving high throughput phenotyping methods are a major opportunity to resolve comparable questions as done here for shoot architecture. We conclude that breeding for specific vigour traits will likely have additional beneficial effects, as indicated by the haplotype studies. Vigour from root growth can then be an added layer at a later step towards field-grown, weed-competitive varieties that can be farmed in a sustainable manner.

## Conclusion

This study explored diversity in shoot architecture of rice seedlings, identified traits contributing to canopy shading potential and identified the putative genetic components related to canopy shading. The traits contributing to a high Shading Rank, and therefore a proposed increased weed-competitiveness, are also intrinsically relevant for seedling vigour. Shoot area, number of leaves and plant height contribute strongly to early vigour and are therefore imperative target traits for weed-competitiveness. We also highlight additional shoot architecture traits, such as solidity and leaf angles, that contribute to increased shading potential and are therefore desirable traits for weed-competitiveness (Figure 2). Indeed, we confirmed that light extinction is significantly stronger under canopies of varieties predicted to have high shading potential and therefore likely being more weed-competitive.

We identified 26 significant marker-trait associations including five novel loci related to canopy shading traits, and the haplotypes corresponding to high-shading potential. Phenotypic investigations carried out in previous studies focused on adult plants and yield traits. This is also reflected in the breeding programme over the last decades, which aimed for high yielding dwarf varieties. Many widely cultivated varieties, such as IR 64 and Nipponbare, showed low Shading Ranks in our analyses, and the most abundant haplotypes, with only few exceptions, were often the ones with lowest shade casting. Our study indicates a clear potential for improvement towards sustainable weed suppression in the current breeding programmes, and that some of the newly studied traits here could be introduced into future breeding programmes.

Summarizing, the acquired knowledge of relevant traits, together with the information about their underlying genomic regions and haplotypes described here can serve as a basis for future reverse-genetic studies and genome-assisted breeding programmes that will contribute to making rice farming more sustainable and help to improve yield in dry, direct-seeded rice.

## Material and methods

### Plant material

344 Asian rice (*O. sativa)* cultivars were used out of an established rice diversity panel (Rice diversity panel 1; RDP1 (Eizenga et al., 2014)). In addition, one African rice variety (*O. glaberrima*) TOG7192 was also included. The RDP1 is a collection of purified, homozygous rice varieties spread over 82 countries all over the world. The panel includes landraces and elite rice cultivars from five subpopulations: *indica* and *aus* belonging to the Indica varietal group and *tropical japonica, temperate japonica* and *aromatic* which comprise the Japonica varietal group, in addition to the *admixture* group, (Liakat Ali et al., 2011; Zhao et al., 2011). The full panel and detailed information (accession name, accession ID, subpopulation and country of origin) can be found in the Supplemental Table 1.

### Growth conditions

Rice plants were grown in the screen-house facilities of the International Rice Research Institute (IRRI)in The Philippines, during October 2017 – April 2018. Temperatures ranged from 37 °C during the day to 27 °C during the night, with a relative humidity of 75 % and 80 %, respectively and a photoperiod ranging from 11 to 12 hours. Four temporally separated replications were carried out, with three plants per variety within each replicated experiment. Plants were grown in a randomized block design in single pots with a 30 cm x 30 cm distance between seedlings. In the first experiment, seeds received from the gene bank at IRRI were exposed to 40 °C for up to 5 days, to break dormancy, followed by 24 h at 21 °C. For germination, seeds were put in Petri dishes (12 per variety) on wet filter paper and incubated at 32 °C for 24 h. Seeds were planted directly on the soil, following the direct-seeded rice method: 4 seeds were placed per pot (diameter of 16 cm and 13 cm high, without drainage holes on the bottom) filled with sterilized clay-loam field soil mixed with complete fertilizer (NPK fertilizer with 46 / 18 / 60 g per kg soil). The seeds were sown at a depth of x-cm and then covered with a thin layer of soil. From planting onwards, soil was kept moist. At 7 days after sowing (DAS), surplus seedlings were removed to retain only 1 seedling per pot. At 14 DAS, fertilizer was added, with 50% of N of concentration off first application. From 15 DAS until the end of the experiment, watering was done to keep a layer of water on the soil and the plants under water-logged conditions.

### Phenotyping

Plants were measured by hand at 28 DAS for the following traits: number of leaves, number of tillers, total plant height, culm height, and length of longest leaf. Plants were photographed from the top and side using 2 digital cameras in a fixed imaging set-up at 21 and 28 DAS. At the last time point, a scan of the blade of the longest leaf was taken and the whole shoot was harvested for analysis of dry weight upon 48 h of drying at 70 °C (IRRI, 2013; Caton et al., 2003). In Table 1, each trait, their abbreviations and evaluation methods are described. The raw data for each replicate can be accessed at https://doi.org/10.5281/zenodo.4730232 (Supplemental Data 1).

### Data processing and statistical analysis

In order to extract traits from RGB images, an automatised image analysis pipeline was established using the open source, python based PlantCV software (PlantCV version 3.7) (Fahlgren et al., 2015; Gehan et al., 2017). We made optimisations to the script for detection of monocots, to enable the extraction of values for shoot area, hull area and perimeter. The python script describing the developed pipeline can be accessed at https://plantcv.readthedocs.io/en/stable/ and the adapted Jupiter notebook used for processing all the images athttps://doi.org/10.5281/zenodo.4730232 (Supplemental Data 2). The measurements of tiller angle, leaf angle and leaf erectness, were done using the free ImageJ software (https://imagej.nih.gov/ij/). Tiller angles were taken between the two outermost tillers and the culm, respectively. The leaf angles were taken between the second and third youngest leaf and the culm, respectively. The leaf droopiness was measured on the same leaves as the interception angle of two tangents aligned to the initiation and the tip of the leaf blade.

The values of the first replicate were excluded for 62 varieties as their position within the greenhouse was more shaded. These positions were excluded from further experimental replication, to ensure equal light conditions for all studied plants. Prior to statistical analysis, the raw data was curated for outliers (using 1.5*IQR away from the mean) and mean was calculated out of the four replicates, with two biological replicates each. Statistical analysis such as ANOVA, Pearson Correlation and Hierarchical Clustering were performed using R (R Version: 3.6.1-1bionic; R Core Team, 2020) and the online tool MVapp https://mvapp.kaust.edu.sa (Julkowska et al., 2019). The Pearson Correlation coefficients between traits were calculated using raw data. For Hierarchical Clustering traits and individual samples were clustered using ward.D2 method. The values of individual traits were normalized per trait using z-Fisher transformation and scaled prior to clustering. Based on the correlation and clustering analysis, a subset of phenotypic traits, was defined as the core traits. The core traits were shoot area, leaf number, solidity, culm height, leaf angle, tiller angle and leaf droopiness. Then we calculated the Shading Rank as follows:

First, we normalized the trait values 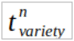

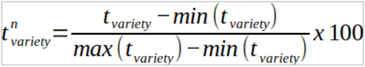

where 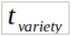 is the value of a certain trait measured for a certain plant in the investigated population and min and max are the minimum and maximum values of the measured trait in the whole population, with the normalized values ranging from 0 to 100.

Next, we calculated the Shading Score for each variety 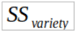

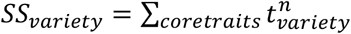 where ∑ is calculated as the sum only from the normalized values of the core traits. From this, we get the Shading Rank (SR), which is the rank given to each variety according to its SS, ordering the varieties from 1 (lowest) to 344 (highest). The list of 344 varieties with their normalized core trait values, the sum of normalized core trait values and their Shading Rank can be found in Supplemental Table 3.

### Canopy shading experiment

Rice were grown in the greenhouse facilities of Utrecht University, in The Netherlands, in February 2021. Temperatures were set to 29 °C during the day and 25 °C during the night and a photoperiod from 8 am to 8 pm, with a minimal light intensity of 400 Чmol m^-2^ s^-1^ and artificial light (Valoya, Model Rx400 500mA 5730, Spectrum AP673L) switching on if sunlight flux rate dropped below 400 Чmol m^-2^ s^-1^. Automatic watering kept soil in pots saturated. The selected *O. sativa* varieties were Shim Balte, Mudgo, Della and Luk Takhar, with Shading Ranks of 344, 330, 49 and 1, respectively. Germination protocol was followed as described above. Four plants were grown per pot, in each of the corners of a square pot (10 x 10 x 11 cm) in a substrate mix of black soil, vermiculite and sand in a ratio of 5 : 3 : 2 together with 6 g Osmocote and 1 l Yoshida nutrient solution per kg substrate. Pots were arranged at a distance of 10 cm in mixed plots. The experiment units (the eight plants that were measured) were surrounded by bordering plants to avoid border effects on the experimental units. Light intensity (photosynthetic active radiation (PAR) of 400-700 nm waveband) was measured at the ground level between rice plants (with six measurements in each of the three replicates) and above the plants for reference at the same time to calculate light extinction. PAR values can be found in Supplemental Table 4.

### Phenotype data for GWAS

For the GWAS analyses, the mean values of all phenotypes were included, only *O. glaberrima* TOG7192 was excluded since it does not belong to the *O. sativa* species. We tested for the normal distribution across the recorded traits prior to running the GWAS. The list for all 344 varieties with 13 shoot trait values (as the mean value out of eight replicates, for raw data see Supplemental Data 1) which were used as input for GWAS can be found in Supplemental Table 5.

### Genotype data

For the genotype data we have used two data sets publicly available at http://ricediversity.org/data/index.cfmtools/. As a second dataset, we used the newer version of genomic data imputed HDRA with 4.8 M SNPs, from 3,010 *O. sativa* varieties assembling the established Rice Reference Panel by merging the high-density rice array with 700 K SNPs from in total 1,568 *O. sativa* varieties including RDP1 (rice diversity panel 1), RDP2 and NIAS (national institute of agrobiological sciences) from (McCouch et al., 2016) and 3000 Rice Genomes data sets (D. R. Wang et al., 2018). The data was curated by filtering for unique SNPs, 90% call rate (90% minimum count) and minor allele frequency ≥ 5 %. We used the SNP data that adhere to the filtering criteria for 344 varieties that were included in the phenotypic screen, which resulted in total of 1.7 M SNPs remained as an input for the GWAS. As an average genome-wide linkage disequilibrium (LD) decay in rice we used previously calculated values (Zhao et al. 2011; Huang et al. 2010). LD is calculated by measuring the pairwise SNP LD among the common SNPs (with MAF > 0.05) using r^2^, the correlation in frequency among pairs of alleles across a pair of markers, using the software PLI NK (http://zzz.bwh.harvard.edu/plink/).

### Genome wide association study (GWAS)

We used two different software packages to perform the GWAS. The first is an R package (R version 3.6.1) of Genomic Association and Prediction Integrated Tool (GAPIT) (Tang et al., 2016; Wang et al., 2018c). We employed a mixed linear model (MLM) (Yu et al., 2006) with the optimal number of Principal Components based on the calculated Bayesian information criterion (BIC) for each trait, including as coefficients a kinship matrix (K-matrix), based on clustering analysis to account for genetic relationship between individuals, together with the population structure (Q-matrix). The Manhattan plots for GWAS using the GAPIT can be found in Supplemental Data 5, for shoot area, hull area, perimeter, plant height, culm height, leaf length, solidity, number of leaves, number of tillers, dry weight, droopiness, leaf angle, tiller angle and the Sum of normalized traits. Shown are SNPs with MAF > 0.05, with the negative logarithmic p-values on the y-axis, for 1.7 M SNPs across the 12 rice chromosomes along the x‐axis. The second software package is lme4QTL (Ziyatdinov et al., 2018). We performed GWAS as described in the paper, taking population structure into account by using a kinship matrix. This kinship matrix was calculated using the cov() function in R 3.6 (Supplemental Figure 2). The decomposition matrix to correct for population structure was made by following the lme4QTL protocol. It uses the relmatLmer(), varcov() and decompose_varcov() functions in order. The obtained decomposition matrix, together with the traits and binary SNP matrix is then used in the matlm() function to calculate the significance and effect per SNP. The full list of detected significant SNP associations can be accessed at https://doi.org/10.5281/zenodo.4730232 (Supplemental Data 3). As a confirmation for the reliability of SNP trait associations, we correlated the results of the two methods applied here (GAPIT and lme4QTL). We do not expect an exact overlap, as there is a small difference in how the kinship matix is calculated and GAPIT uses MLM, whereas lme4QTL does not. The narrow sense heritability (*h*^*2*^) of the analysed traits was calculated with GAPIT (Supplemental Table 6). To set the significance threshold the rather conservative Bonferroni correction was applied, calculated by the –log^10^(p-value of 0.05/ΣSNPs), which corresponds to −log10(0.05/1.700.000) = −7.53 for the imputed HDRA data set. To examine the GWAS model performance and estimate possible model overfitting, QQ plots were generated (Supplemental Data 6).

### Post-GWAS analysis

For all follow-up analysis the output of the GWAS on the raw, untransformed phenotype data was used.

#### Locus definition

We determined loci to be of interest, if there are several significantly associated SNPs found in close proximity. Single SNPs passing the threshold were neglected, because whole-genome sequencing data provides enough markers in each linkage disequilibrium block. Since rice has a low rate of LD decay, this makes it more difficult to identify causal genes (Wang et al., 2020). Therefore, the local LD analysis was used to define LD clumps surrounding the index SNPs, using LD clumping in PLINK, where the local LD between SNPs is considered. A strong LD between SNPs is one of the three criteria that must be simultaneously satisfied. The other two criteria are p-value threshold set to 0.01 and physical distance set to 250 kb, given with the R^2^ value. We considered SNPs with –log^10^(p-value) > 5 as index SNPs to perform the analysis and clump SNPs with p-value > 4. For the determination of loci of interest for weed-competitiveness, we focused on the core traits culm height, shoot area, solidity and number of leaves. For culm height and number of leaves single significant SNPs were not found to be surrounded by other significant SNPs within LD and therefore did not meet our selection cr**i**teria. Since, dry weight is highly correlated with the traits of branchiness, we included the peaks found for dry weight as a representative locus for branchiness and similarly the loci for plant height as a representative of height related traits.

#### Gene models

Genetic regions covered by significant SNPs were searched for candidate genes using two different gene annotation models, which were then merged: the Michigan State University (MSU; 31 Oct. 2011 - Release 7; http://rice.plantbiology.msu.edu/) and the Rice Annotation Project Database (RAP-DB; 24 March 2020; https://rapdb.dna.affrc.go.jp/). Other data resources used, were the gene ID converter (https://rapdb.dna.affrc.go.jp/tools/converter), GALAXY – rice genome browser (http://13.250.174.27:8080/?tool_id=getgenes&version=1.0.0&identifer=pxuu9t4bnk) and SNP seek (http://snp-seek.irri.org/).

### Haplotype analysis

In order to facilitate the identification of candidate genes within the found loci related to the canopy architecture, we performed haplotype analysis spanning the coding sequence regions of the genes within each locus. For each locus, we used the combined gene model annotation (MSU and RAP-DB) to identify the coding sequences belonging to individual genes (Supplemental Table 7). We subsequently compiled all SNPs that were within the coding sequence region into one haplotype and grouped all studied varieties based on their haplotype sequence. The haplotypes represented by 2 or less varieties were excluded from the analysis, due to low representation. Based on the haplotype grouping for each coding sequence, we performed a t-test for significant differences between the most abundant haplotype with all the other identified haplotypes for all measured traits. The individual haplotypes are represented by A/T, where A stands for reference accession sequence, and T for any alternative variant. Supplemental Data 8 contains the full list of coding sequences of genes within the defined loci of interest.

## Acknowledgements

We thank Ricardo Eugenio and James Edgane for their substantial assistance in the phenotyping at the International Rice Research Institute and Yorrit van de Kaa and Alba Schielen for their help with the experiments at Utrecht University. We thank Roel van Bezouw and Tom Theeuwen for helpful discussions about GWAS and Rens Voesenek, Evelyn Aparicio (Nelen & Schuurmans), Jochem Evers (WUR) and Jonne Rodenburg (University of Greenwich) for useful discussions on this research project.

## Supplemental Data

**Supplemental Table 1:** List of rice varieties of screened rice diversity panel (RDP1) and description of origin.

**Supplemental Table 2:** Results for ANOVA (considered significant with p < 0.05) and post-hoc based on Tukey’s pairwise comparison of shoot traits between different rice varieties and between different subpopulations, mean out of eight replicates of 344 varieties, the sum of normalized core trait values and their Shading Rank. Raw data can be found in https://doi.org/10.5281/zenodo.4730232 (Supplemental Data 1).

**Supplemental Table 3:** The list of 344 varieties with their normalized core trait values, the sum of normalized core trait values and their Shading Rank.

**Supplemental Table 4:** PAR values (photosynthetic active radiation of 400-700 nm waveband) and measured measured at the ground level under the rice canopy and reduction in light intensity (% PAR) compared to above the canopy for different rice varieties.

**Supplemental Table 5:** List of 344 varieties with 13 shoot trait values (as the mean value out of eight replicates, for raw data see Supplemental Data 1) which were used as input for genome-wide association studies, their normalized trait values, the sum of normalized core trait values and their Shading Rank.

**Supplemental Table 6:** Narrow sense heritability of all analysed traits in genome-wide association studies, calculated in GAPIT.

S**upplemental Table 7:** Full list of SNP positions in loci of interest with gene annotation and gene ontology categories from Rice Annotation Project Database.

**Supplemental Figure 1:** Scatter plots and R^2^ values for pair-wise correlation analysis for individual traits.

**Supplemental Figure 2:** Kinship matrix of screened rice diversity panel (RDP1).

**Supplemental Data 1** (https://doi.org/10.5281/zenodo.4730232): List of 344 varieties with raw data of 13 shoot traits from eight replicates.

**Supplemental Data 2** (https://doi.org/10.5281/zenodo.4730232): Python script based on PlantCV used for image analysis.

**Supplemental Data 3** (https://doi.org/10.5281/zenodo.4730232): Association results for GWAS with Lme4QTL using a mixed linear model (MLM) based on the lme4QTL protocol, for shoot area, hull area, perimeter, plant height, culm height, leaf length, solidity, number of leaves, number of tillers, dry weight, droopiness, leaf angle, tiller angle and the Sum of normalized traits.

**Supplemental Data 4:** Genetic regions underlying shoot architectural traits and seedling vigour in 4-week-old rice seedlings. Single-trait genome-wide association studies (GWAS) using a mixed linear model (MLM) based on the lme4QTL protocol, for droopiness, leaf angle, tiller angle, SUM_norm_traits, number of leaves, number of tillers, culm height, leaf length hull area and perimeter. The Manhattan plots depict the single nucleotide polymorphisms (SNPs) with minor allele frequencies (MAF) > 0.05. Negative logarithmic P-values on the y-axis, for 1.7 M SNPs across the 12 rice chromosomes along the x‐axis. P-values of association results for all traits can be found in Supplemental Data 3.

**Supplemental Data 5:** Genetic regions underlying shoot architectural traits and seedling vigour in 4-week-old rice seedlings. Single-trait GWAS using a mixed linear model (MLM) with the GAPIT package in R, for shoot area, hull area, perimeter, plant height, culm height, leaf length, solidity, number of leaves, number of tillers, dry weight, droopiness, leaf angle, tiller angle and the Sum of normalized traits. The Manhattan plots depict the single nucleotide polymorphisms (SNPs) with minor allele frequencies (MAF) > 0.05. Negative logarithmic P values on the y-axis, for 1.7 M SNPs across the 12 rice chromosomes along the x‐axis.

**Supplemental Data 6:** QQ plots with negative logarithmic P values for observed on the y-axis and expected SNP - trait associations on the x-axis.

**Supplemental Data 7:** Haplotype groups for all determined loci of interest with their phenotype effect for 13 investigated shoot traits.

**Supplemental Data 8:** List of sequences of genes for loci of interest, with haplotypes for screened varieties.

## References

Acevedo‐Siaca, L.G., Coe, R., Wang, Y., Kromdijk, J., Quick, W.P., and Long, S.P. (2020). Variation in photosynthetic induction between rice accessions and its potential for improving productivity. New Phytol. nph.16454.

Andrew, I.K.S., Storkey, J., and Sparkes, D.L. (2015). A review of the potential for competitive cereal cultivars as a tool in integrated weed management. Weed Res. 55, 239–248.

Ballaré, C.L., and Pierik, R. (2017). The shade-avoidance syndrome: multiple signals and ecological consequences. Plant. Cell Environ. 40, 2530–2543.

Brainard, D.C., Bellinder, R.R., and DiTommaso, A. (2005). Effects of canopy shade on the morphology, phenology, and seed characteristics of Powell amaranth (Amaranthus powellii). Weed Sci. 53, 175–186.

Casal, J.J. (2012). Shade Avoidance. 1–19.

Caton, B.P., Cope, A.E., and Mortimer, M. (2003). Growth traits of diverse rice cultivars under severe competition: Implications for screening for competitiveness. F. Crop. Res. 83, 157–172.

Chakraborty, D., Ladha, J.K., Rana, D.S., Jat, M.L., Gathala, M.K., Yadav, S., Rao, A.N., Ramesha, M.S., and Raman, A. (2017). A global analysis of alternative tillage and crop establishment practices for economically and environmentally efficient rice production. Sci. Rep. 7, 1–11.

Chauhan, B.S. (2012). Weed Ecology and Weed Management Strategies for Dry-Seeded Rice in Asia. Weed Technol. 26, 1–13.

Chauhan, B., and Yadav, A. (2013). Weed management approaches for dry-seeded rice in India: a review. Indian J. Weed Sci. 45, 1–6.

Chauhan, B.S., Jabran, K., and Mahajan, G. (2017). Rice Production Worldwide (Springer Nature).

Chen, K., Zhang, Q., Wang, C.C., Liu, Z.X., Jiang, Y.J., Zhai, L.Y., Zheng, T.Q., Xu, J.L., and Li, Z.K. (2019). Genetic dissection of seedling vigour in a diverse panel from the 3,000 Rice (Oryza sativa L.) Genome Project. Sci. Rep. 9.

Cordero-Lara, K.I., Kim, H., and Tai, T.H. (2016). Identification of Seedling Vigor-Associated Quantitative Trait Loci in Temperate Japonica Rice. Plant Breed. Biotechnol. 4, 426–440.

Dimaano, N.G.B., Ali, J., Sta. Cruz, P.C., Baltazar, A.M., Diaz, M.G.Q., Acero, B.L., and Li, Z. (2017). Performance of Newly Developed Weed-Competitive Rice Cultivars under Lowland and Upland Weedy Conditions. Weed Sci. 65, 798–817.

Dingkuhn, M., Tivet, F., Siband, P.-L., Asch, F., Audebert, A., Sow, A., and International Rice Research Conference. Los Banos Philippines), P. 2000-03-31/2000-04-03; I. (Los B. (2001). Varietal differences in specific leaf area: a common physiological determinant of tillering ability and early growth vigor? In Rice Research for Food Security and Poverty Alleviation. Proceedings of the International Rice Research Conference, 31 March - 3 April 2000, S. Peng, and B. Hardy, eds. (Los Banos, Philippines), pp. 95–108.

Eizenga, G.C., Ali, M.L., Bryant, R.J., Yeater, K.M., McClung, A.M., and McCouch, S.R. (2014). Registration of the Rice Diversity Panel 1 for Genomewide Association Studies. J. Plant Regist. 8, 109.

Fahlgren, N., Feldman, M., Gehan, M.A., Wilson, M.S., Shyu, C., Bryant, D.W., Hill, S.T., McEntee, C.J., Warnasooriya, S.N., Kumar, I., et al. (2015). A versatile phenotyping system and analytics platform reveals diverse temporal responses to water availability in Setaria. Mol. Plant 8, 1520–1535.

Fao, F. and A.O. of the U.N. (2019). World Food and Agriculture – Statistical pocketbook 2019 (Rome).

Farooq, M., Siddique, K.H.M.M., Rehman, H., Aziz, T., Lee, D.-J.J., and Wahid, A. (2011). Rice direct seeding: Experiences, challenges and opportunities.

Franklin, K.A. (2008). Shade avoidance. New Phytol. 179, 930–944.

Gehan, M.A., Fahlgren, N., Abbasi, A., Berry, J.C., Callen, S.T., Chavez, L., Doust, A.N., Feldman, M.J., Gilbert, K.B., Hodge, J.G., et al. (2017). PlantCV v2: Image analysis software for high-throughput plant phenotyping. PeerJ 5, e4088.

Ghosal, S., Casal, C., Quilloy, F.A., Septiningsih, E.M., Mendioro, M.S., and Dixit, S. (2019). Deciphering Genetics Underlying Stable Anaerobic Germination in Rice: Phenotyping, QTL Identification, and Interaction Analysis. Rice 12.

Haefele, S. M., Johnson, D. E., M’Bodj, D., Wopereis, M.C. C.S., Miezan, K. M., M’Bodj, D., Wopereis, M.C. C.S., and Miezan, K. M. (2004). Field screening of diverse rice genotypes for weed competitiveness in irrigated lowland ecosystems. F. Crop. Res. 88, 29–46.

Huang, X., Wei, X., Sang, T., Zhao, Q., Feng, Q., Zhao, Y., Li, C., Zhu, C., Lu, T., Zhang, Z., et al. (2010). Genome-wide association studies of 14 agronomic traits in rice landraces. Nat. Genet. 42, 961–967.

International Rice Research Institute (2013). SES (Standard Evaluation System) for Rice (Manila, Philippines).

Julkowska, M.M., Saade, S., Agarwal, G., Gao, G., Pailles, Y., Morton, M., Awlia, M., and Tester, M. (2019). MVApp—Multivariate Analysis Application for Streamlined Data Analysis and Curation. 180, 1261–1276.

Kennedy, G., and Burlingame, B. (2003). Analysis of food composition data on rice from a plant genetic resources perspective. Food Chemistry 80, 589–596.

Kretzschmar, T., Pelayo, M.A.F., Trijatmiko, K.R., Gabunada, L.F.M., Alam, R., Jimenez, R., Mendioro, M.S., Slamet-Loedin, I.H., Sreenivasulu, N., Bailey-Serres, J., et al. (2015). A trehalose-6-phosphate phosphatase enhances anaerobic germination tolerance in rice. Nat. Plants 1, 1–5.

Kumar, V., and Ladha, J.K. (2011). Direct Seeding of Rice. Recent Developments and Future Research Needs (Academic Press).

Li, M., Liu, X., Bradbury, P., Yu, J., Zhang, Y.M., Todhunter, R.J., Buckler, E.S., and Zhang, Z. (2014). Enrichment of statistical power for genome-wide association studies. BMC Biol. 12, 73.

Liakat Ali, M., McClung, A.M., Jia, M.H., Kimball, J.A.J.A., McCouch, S.R., Eizenga, G.C., McCouch, S.R., and Georgia, C.E. (2011). A Rice Diversity Panel Evaluated for Genetic and Agro-Morphological Diversity between Subpopulations and its Geographic Distribution. Crop Sci. 51, 2021–2035.

Mackill, D.J., and Khush, G.S. (2018). IR64: a high-quality and high-yielding mega variety. Rice 11, 18.

Mahajan, G., and Chauhan, B.S. (2013). The role of cultivars in managing weeds in dry-seeded rice production systems. Crop Prot.

McCouch, S.R., Wright, M.H., Tung, C.-W.W., Maron, L.G., McNally, K.L., Fitzgerald, M., Singh, N., DeClerck, G., Agosto-Perez, F., Korniliev, P., et al. (2016). Open access resources for genome-wide association mapping in rice. Nat. Commun. 7, 10532.

Mennan, H., Ngouajio, M., Sahin, M., Isık, D., and Altop, K. (2012). Competitiveness of rice (Oryza sativa L.) cultivars against Echinochloa crus-galli (L.) Beauv. in water-seeded production systems. Crop Prot. 41, 1– 9.

Namuco, O.S.S., Cairns, J.E.E., and Johnson, D.E.E. (2009). Investigating early vigour in upland rice (Oryza sativa L.): Part I. Seedling growth and grain yield in competition with weeds. F. Crop. Res. 113, 197–206.

Oliver, V., Cochrane, N., Magnusson, J., Brachi, E., Monaco, S., Volante, A., Courtois, B., Vale, G., Price, A., and Teh, Y.A. (2019). Effects of water management and cultivar on carbon dynamics, plant productivity and biomass allocation in European rice systems. Sci. Total Environ. 685, 1139–1151.

Pierik, R., and De Wit, M. (2014). Shade avoidance: phytochrome signalling and other aboveground neighbour detection cues. J. Exp. Bot. 65, 2815–2824.

Poorter, H., Niklas, K.J., Reich, P.B., Oleksyn, J., Poot, P., and Mommer, L. (2012). Biomass allocation to leaves, stems and roots: meta-analyses of interspecific variation and environmental control. New Phytol. 193, 30–50.

R Core Team (2020). R: A language and environment for statistical computing. R Foundation for Statistical Computing, Vienna, Austria. URL https://www.R-project.org/.

Rao, A.N., Johnson, D.E., Sivaprasad, B., Ladha, J.K., and Mortimer, A.M. (2007). Weed Management in Direct-Seeded Rice. Adv. Agron. 93, 153–255.

Roig-Villanova, I., and Martínez-García, J.F. (2016). Plant Responses to Vegetation Proximity: A Whole Life Avoiding Shade. Front. Plant Sci. 7, 236.

Sakamoto, T., Morinaka, Y., Ohnishi, T., Sunohara, H., Fujioka, S., Ueguchi-Tanaka, M., Mizutani, M., Sakata, K., Takatsuto, S., Yoshida, S., et al. (2006). Erect leaves caused by brassinosteroid deficiency increase biomass production and grain yield in rice. Nat. Biotechnol. 24, 105–109.

Seavers, G.P., and Wright, K.J. (1999). Crop canopy development and structure influence weed suppression. Weed Res. 39, 319–328.

Subedi, S.R., Sandhu, N., Singh, V.K., Sinha, P., Kumar, S., Singh, S.P., Ghimire, S.K., Pandey, M., Yadaw, R.B., Varshney, R.K., et al. (2019). Genome-wide association study reveals significant genomic regions for improving yield, adaptability of rice under dry direct seeded cultivation condition. BMC Genomics 20, 471.

Tang, Y., Liu, X., Wang, J., Li, M., Wang, Q., Tian, F., Su, Z., Pan, Y., Liu, D., Lipka, A.E., et al. (2016). GAPIT Version 2: An Enhanced Integrated Tool for Genomic Association and Prediction. Plant Genome 9.

Teichmann, T., and Muhr, M. (2015). Shaping plant architecture. Front. Plant Sci. 6, 233.

Wang, D.R., Agosto-Pérez, F.J., Chebotarov, D., Shi, Y., Marchini, J., Fitzgerald, M., McNally, K.L., Alexandrov, N., and McCouch, S.R. (2018a). An imputation platform to enhance integration of rice genetic resources. Nat. Commun. 9, 3519.

Wang, F., Longkumer, T., Catausan, S.C., Calumpang, C.L.F., Tarun, J.A., Cattin-Ortola, J., Ishizaki, T., Pariasca Tanaka, J., Rose, T., Wissuwa, M., et al. (2018b). Genome-wide association and gene validation studies for early root vigour to improve direct seeding of rice. Plant Cell Environ. 41, 2731–2743.

Wang, Q., Tian, F., Pan, Y., Buckler, E.S., and Zhang, Z. (2018c). User Manual for Genomic Association and Prediction Integrated Tool.

Wang, Q., Tang, J., Han, B., and Huang, X. (2020). Advances in genome-wide association studies of complex traits in rice. Theor. Appl. Genet. 1415–1425.

Weiner, J., Andersen, S.B., Wille, W.K., Griepentrog, H.W., and Olsen, J.M. (2010). Evolutionary Agroecology: the potential for cooperative, high density, weed-suppressing cereals. Evol. Appl. 3, 473–479.

Wing, R.A., Purugganan, M.D., and Zhang, Q. (2018). The rice genome revolution: from an ancient grain to Green Super Rice. Nat. Rev. Genet. 1.

Worthington, M., and Reberg-Horton, C. (2013). Breeding Cereal Crops for Enhanced Weed Suppression: Optimizing Allelopathy and Competitive Ability. J. Chem. Ecol. 39, 213–231.

Xu, L., Li, X., Wang, X., Xiong, D., and Wang, F. (2019). Comparing the grain yields of direct-seeded and transplanted rice: A meta-analysis. Agronomy 9, 767.

Yang, W., Guo, Z., Huang, C., Duan, L., Chen, G., Jiang, N., Fang, W., Feng, H., Xie, W., Lian, X., et al. (2014). Combining high-throughput phenotyping and genome-wide association studies to reveal natural genetic variation in rice. Nat. Commun. 5.

Yu, J., Pressoir, G., Briggs, W.H., Bi, I.V., Yamasaki, M., Doebley, J.F., McMullen, M.D., Gaut, B.S., Nielsen, D.M., Holland, J.B., et al. (2006). A unified mixed-model method for association mapping that accounts for multiple levels of relatedness. Nat. Genet. 38, 203–208.

Zhang, P., Kowalchuk, G.A., Soons, M.B., Hefting, M.M., Chu, C., Firn, J., Brown, C.S., Zhou, X.X., Zhou, X.X., Guo, Z., et al. (2019). SRU D: A simple non‐destructive method for accurate quantification of plant diversity dynamics. J. Ecol. 107, 2155–2166.

Zhang, Z.-H., Wang, K., Guo, L., Zhu, Y.-J., Fan, Y.-Y., Cheng, S.-H., and Zhuang, J.-Y. (2012). Pleiotropism of the Photoperiod-Insensitive Allele of Hd1 on Heading Date, Plant Height and Yield Traits in Rice. PLoS One 7, e52538.

Zhao, D.L., Atlin, G.N., Bastiaans, L., and Spiertz, J.H.J. (2006a). Developing selection protocols for weed competitiveness in aerobic rice. F. Crop. Res. 97, 272–285.

Zhao, D.L., Atlin, G.N., Bastiaans, L., and Spiertz, J.H.J. (2006b). Comparing rice germplasm groups for growth, grain yield and weed-suppressive ability under aerobic soil conditions. Weed Res. 46, 444–452.

Zhao, D.L., Bastiaans, L., Atlin, G.N., and Spiertz, J.H.J. (2007). Interaction of genotype × management on vegetative growth and weed suppression of aerobic rice. F. Crop. Res. 100, 327–340.

Zhao, K., Tung, C., Eizenga, G.C., Wright, M.H., Ali, M.L., Price, A.H., Norton, G.J., Islam, M.R., Reynolds, A., Mezey, J., et al. (2011). Genome-wide association mapping reveals a rich genetic architecture of complex traits in Oryza sativa. Nat. Commun. 2, 1–10.

Ziyatdinov, A., Vázquez-Santiago, M., Brunel, H., Martinez-Perez, A., Aschard, H., and Soria, J.M. (2018). lme4qtl: linear mixed models with flexible covariance structure for genetic studies of related individuals. BMC Bioinformatics 19, 68.

